# Design-informed Size Factor Estimation

**DOI:** 10.64898/2026.08.13.744630

**Authors:** Todd Pocuca, Guillaume Paré, Benjamin M. Bolker

## Abstract

Accurate normalization is essential for differential expression analysis of RNA-sequencing data. Popular normalization methods such as the median-of-ratios and trimmed mean of M-values do not leverage information from the experimental design. This may be inefficient in experiments with large-scale systematic expression changes or complex designs. Here, we introduce design-informed size factor estimation (disize), a normalization method that uses information from the experimental design to improve accuracy. disize uses a modified generalized linear mixed model to robustly distinguish between biological signal and sample-specific size factors. We also propose a mechanistically justified data-generating process for RNA-sequencing counts that is derived from previous models of transcription and sequencing. Through simulations based on this data-generating process and validating on true RNA-seq data, we show that disize recovers size factors more accurately than existing methods, particularly in challenging scenarios with low gene expression and a high proportion of differentially expressed genes; this in turn improves downstream analysis. disize provides a robust and accurate approach to normalization, highlighting the significant benefits of integrating experimental design information directly into normalization for transcriptomic datasets.

**Author summary:** In transcriptomic analysis, normalization adjusts for technical biases arising from library preparation and sequencing. Methods implemented in widely used packages like DESeq2 and edgeR ignore information in the experimental design during normalization.

Incorporating information from the experimental design into a normalization method has the potential to yield more accurate results. To do this, we developed a new method, design-informed size factor estimation (disize), that uses a statistical model to jointly account for the biological signal defined by the design and the sample-specific batch effect. By separating the biological variation into its components, disize can more robustly estimate the batch effect. To validate our approach, we constructed a flexible simulation framework relying on a mechanistically justified data-generating process for RNA-seq data. Our benchmarks on both simulated and true RNA-seq data show that disize recovers the true size factors more accurately than existing methods, particularly in challenging scenarios with low counts or a high proportion of differentially expressed genes. This improved normalization yields more reliable downstream results in differential expression analysis.

## Introduction

The falling costs of high-throughput sequencing over the last decade have led to an increase in the number of transcriptomic experiments, with applications in various fields ranging from oncology [1] to ecology [2]. This surge in data has in turn driven the development of statistical models to analyze the resulting RNA sequencing (RNA-seq) data. Among the most widely used tools for RNA-seq analysis are DESeq [3], its successor DESeq2 [4], and edgeR [5], all of which provide flexible frameworks for quantifying differential expression (DE).

Although DESeq and edgeR have evolved considerably since their inception, this progress has largely focused on refining downstream DE testing methodology. However, the accuracy of DE analysis fundamentally depends on proper normalization, as technical biases from factors like library preparation, sequencing depth, and RNA composition can distort results [6]. To address these biases, both tools rely on default normalization methods—the median-of-ratios (MoR) [3] in DESeq/DESeq2 and the trimmed mean of M-values (TMM) [6] in edgeR—that assume most genes are not differentially expressed across samples [3, 6]. However, this assumption may sometimes be violated strongly enough to compromise the accuracy of these methods.

Furthermore, a shared characteristic of these methods is that the information from the experimental design is considered only during DE testing, not during normalization. Including design information may improve normalization accuracy, particularly when there is systematic biological variation between groups of samples. For example, samples belonging to the same experimental unit are likely to have similar expression profiles, while samples from different units often have systematic differences. This suggests that normalization could be improved by accounting for the biological variation between samples due to the experimental design. Motivated by this, we propose **d**esign-**i**nformed **size** factor estimation (disize), which explicitly models both biology and batch effects by directly incorporating the experimental design during estimation. We evaluate our method in both bulk and single-cell workflows to show its robustness across bulk and pseudo-bulk contexts.

It is also crucial to distinguish between how well a given normalization method estimates the technical bias (quantified by an estimated *size factor*) and its effects on the downstream DE analysis. An ideal normalization method would recover these latent technical factors perfectly, thereby providing an upper bound for performance; if we knew the true size factors, it should be impossible to perform better in *any* downstream DE analysis. Previous benchmarks have focused exclusively on the effects of normalization on DE testing, which, while important, fails to acknowledge that normalization can only improve a downstream analysis if the estimated size factors accurately recover the underlying technical biases. This oversight may be in part due to the lack of a detailed data-generating process (DGP) for transcriptomic experiments, which should follow from our mechanistic understanding of transcription and the measurement probe. Therefore, we additionally suggest a mechanistically justified DGP for RNA-seq experiments that explicitly models the effect of technical factors (batch effects) on the measured count distribution.

## Materials and methods

### The Data-Generating Process of RNA-Seq Data

We begin the construction of the DGP for RNA-seq data by considering the steady-state distribution of transcript counts of single cells within a homogeneous cell population. Let *c ∈ C* index a cell, *g ∈ G* index a gene, and *X_c,g_* denote the number of transcripts of gene *g* for cell *c*. We assign a negative binomial (NB) distribution to *X_c,g_* based on the model of transcription examined in [7], where the steady-state distribution of transcript counts is well described by a NB fit:

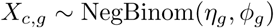

We use *η_g_*to denote the expected count (a measure of the magnitude of expression) and *ϕ_g_* to denote the (inverse) overdispersion factor (a measure of variability):

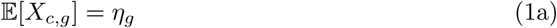

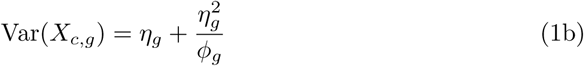

However, in bulk experiments (or single-cell experiments where pseudo-bulking is performed) transcripts from these cells are pooled together, which implies the number of transcripts for a given sample *X_g_*=Σ*_cɛC_ X_c,g_* is distributed according to a NB with scaled parameters:

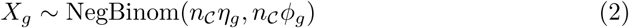

where *n_C_* = *|C|*. With the use of unique molecular identifiers (UMIs), the effect of library preparation and sequencing on the measured count has previously been modelled by a binomial thinning operation [8] on the original number of transcript counts [9]. In other words, given *X_g_*, the total technical effect *ρ* stochastically scales down the original number of transcripts:

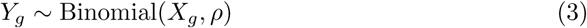

However, since *X_g_*is never observed, the compound distribution produced as a result of marginalizing over *X_g_*is given by:

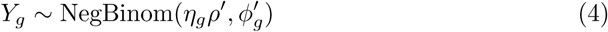

where *ρ^′^* has absorbed the *n_C_* term (as it is a sample-specific quantity), and *ϕ_g_^′^* = *n_C_ϕ_g_*. Thus, the proposed DGP implies the measured count distribution can be described by a NB with parameters related to the original biological signal *η_g_^′^, ϕ_g_^′^* and a sample-specific technical bias *ρ^′^* that scales this distribution.

It is worth noting that the process above assumes homogeneity within the population *C*, which is only true in pure cell cultures. In tissue-level bulk sequencing, the measured count is an aggregation of a heterogeneous mixture of distinct cell types, but the sum of negative binomial random variables with varying parameters does not itself yield a negative binomial. However, the resulting distribution is in practice well-approximated by a negative binomial. We deliberately adopt this simplified homogeneous framing to clearly demonstrate how a measured count can be cleanly decomposed into its underlying biological expression and the batch effect without loss of generality.

### Implementation of disize

disize uses the L-BFGS algorithm offered in the probabilistic programming language Stan to find the maximum a posteriori of a Bayesian model that jointly estimates the effect of covariates (structured according to the experimental design) on gene expression and the batch effect.

The model can be described as a modification of a simpler generalized linear mixed model with offsets:

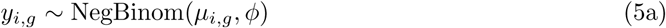

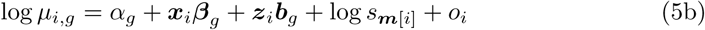

where *y_i,g_* denotes the observed count for a sample *i* and gene *g*. The count is realized from a negative binomial with a mean described by the effect of the covariates *α_g_* + ***x****_i_****β****_g_* + ***z****_i_****b****_g_* (structured according to the experimental design), any specified offset *o_i_* adjusting for known variation, the batch effect *s****_m_***_[*i*]_, structured according to the batch membership ***m***, and a shared inverse overdispersion factor *ϕ*. Here, the inverse overdispersion factor *ϕ* is completely pooled across genes rather than estimated per-gene (*ϕ_g_*) as an approximation to reduce model complexity.

Since the size factors ***s*** are unobserved (and thus different from true offsets), this model is not identifiable for most experimental designs: we cannot distinguish between batch and biology. This problem is overcome by constraining ***s*** to sum to the total number of batches, and assuming only a subset of features are affected by the covariates measured in the experiment. In other words, the estimated coefficients ***β****_g_,* ***b****_g_* (excluding the intercept) are *sparse* across genes.

We encode this assumption by placing separate horseshoe priors [10] on each of the model coefficients. This structure allows different degrees of sparsity for each predictor:

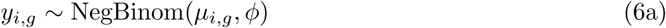

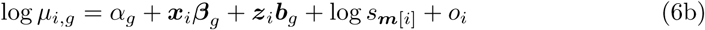

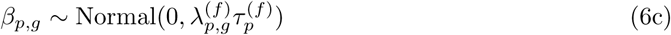

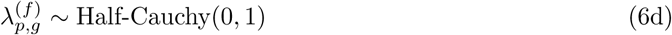

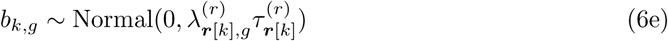

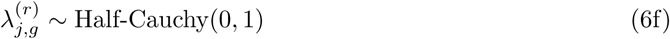

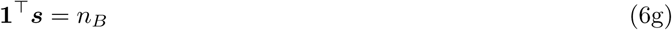

where *p* indexes the fixed-effect coefficients, *k* indexes the random-effect coefficients, *j* indexes distinct random-effect terms, and ***r*** denotes the term membership structure for the random-effect coefficients.

The Stan implementation splits the data and parameter matrices into chunks gene-wise in order to evaluate all distribution statements in parallel. The source code for disize is available at https://github.com/toddpocuca/disize, archived at Zenodo [11].”

### Simulation Framework

The suggested DGP above implies the expectation of a count matrix can be decomposed into the effects of the experimental design on the true biological signal and the batch effect. Therefore, we can construct a count matrix ***Y*** with *n_i_* observations (samples) and *n_g_* features (genes) for a specified experimental design (which defines the fixed-and random-effects model matrices, ***X*** and ***Z***, respectively) and the following parameters:

1. n_genes (*n_g_*): The number of genes to simulate.
2. sparsity (*p*): The fraction of genes that are *not* affected by the experimental conditions (i.e., the non-differentially expressed genes).
3. mgt (*σ*): How strongly the experimental design affects genes that are DE.
4. avg (*δ*): The average baseline expression.

For each batch *b*, we draw an absolute size factor *ρ_b_* (analogous to the technical effect in Eq 3) from a uniform distribution, *ρ_b_ ∼* Uniform(0.1, 1.0). For each gene *g*, we next draw an inverse overdispersion factor *ϕ_g_* from a log-normal distribution, *ϕ_g_ ∼* LogNormal(log 10, 0.5), to mimic the variance in overdispersion present in realistic count data. A gene-specific baseline expression *α_g_* is drawn from a normal distribution with mean log *δ*and standard deviation 1. Each fixed-effect coefficient *β_p,g_* and block of random-effect coefficients *β_k,g_* (belonging to the same term) is then sampled from a mixture of a point mass at 0 with probability *p*, and a centered normal distribution with scale *σ* and probability 1 *− p*. The log of the expected count, log *µ_i,g_*, is then calculated as the sum of the baseline expression, effect from the experimental design, and the log-transformed absolute size factor:

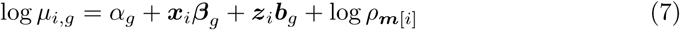

where ***x****_i_* and ***z****_i_* are rows from the fixed-and random-effects model matrices defined by the experimental design, and ***m*** specifies the batch membership structure for each sample. Finally, the observed count for a sample *i* and gene *g* is drawn from a NB with parameters *µ_i,g_* and *ϕ_g_*:

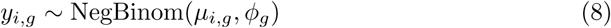

### Generation of Ground-truth Pseudo-bulk RNA-seq Data

While parametric simulations allow for the generation of synthetic RNA-seq counts without existing data, they may fail to capture biological variation and technical noise inherent to true sequencing experiments. Conversely, transcriptomic datasets featuring both complex experimental designs and ground-truth size factors are exceedingly rare. To bridge this gap, we use single-sample single-cell RNA sequencing (scRNA-seq) datasets to construct pseudo-bulk RNA-seq data with pre-defined size factors through a simulated pooling strategy. By artificially partitioning cells from the same sample into distinct groups, we can mimic multi-batch experimental layouts while preserving any unspecified variation in the original data. Similar frameworks using binomial thinning have previously been implemented to evaluate differential expression testing in both bulk and single-cell workflows [12].

Given a scRNA-seq count matrix of a single sample, cells within each annotated cell-type cluster are randomly assigned to one of *n_B_* batches. For each distinct combination of cluster *k* and batch *b*, a pseudo-bulk expression profile is generated by summing the counts across all assigned cells. Because the number of cells randomly assigned to each batch-cluster combination may vary, the overall library size across batches within each cluster must be standardized. Let ***y****_k,b_* represent the raw pseudo-bulk count vector for cluster *k* in batch *b*, and let *N_k,b_* denote the total number of cells contributing to that profile. We compute a balancing weight *w_k,b_* as:

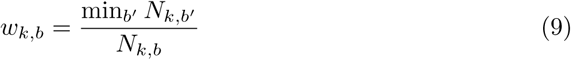

An intermediate, balanced count for gene *g* is then generated through binomial thinning:

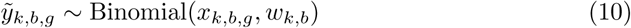

This constructs an intermediate pseudo-bulk dataset with no batch effects present across groups. Finally, user-defined ground-truth size factors ***s*** = [*s*_1_*, s*_2_*,…,s_B_*] are imposed to simulate batch effects:

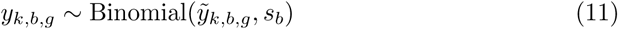

The advantage of this two-stage binomial thinning process is that all unprespecified biological variation inherent to the source tissue is implicitly maintained.

## Results

### Size Factor Estimation Accuracy

We evaluated the performance of disize against established normalization methods (MoR and TMM) across three distinct experimental designs, evaluating each parameter setting across 100 independent simulation replicates (*n*_sims_ = 100) simulating *n_g_* = 10, 000 genes. Each set of simulations varied the baseline expression levels (avg, *δ*) from an average of *δ* = 10.0 to *δ* = 1000.0, evaluated the proportion of non-differentially expressed genes (sparsity, *p*) at low *p* = 20% and high *p* = 80% levels of sparsity, and fixed the magnitude of covariate effects (mgt, *σ*) to capture both weak *σ* = 0.5 and strong *σ* = 2.0 responses to the experiment. Specifically, we analyzed a trivial baseline design with non-systematic variation and no active condition contrasts across 8 donors (a random-intercept-only design), a standard two-condition design with 10 total donors split equally between two groups (5 replicates per condition), and a multi-factorial design crossing two conditions and sex in a 2 *×* 2 factorial structure using 12 total donors (3 replicates per experimental unit).

In the trivial experimental scenario with no systematic variation, size factor estimation accuracy was largely comparable across all three methods (S1 Fig). However, in experiments featuring systematic biological variation, disize showed a clear advantage, proving robust to low sparsity and large biological variation. Specifically, in low-sparsity environments where the majority of genes (80%, *p* = 0.2) were truly differentially expressed, MoR and TMM estimates degraded significantly relative to disize even with a high average baseline expression level (Fig 1, S2 and S3 Figs). This advantage was most pronounced in challenging settings with low average baseline expression levels (*<* 50 counts).

**Fig 1.**
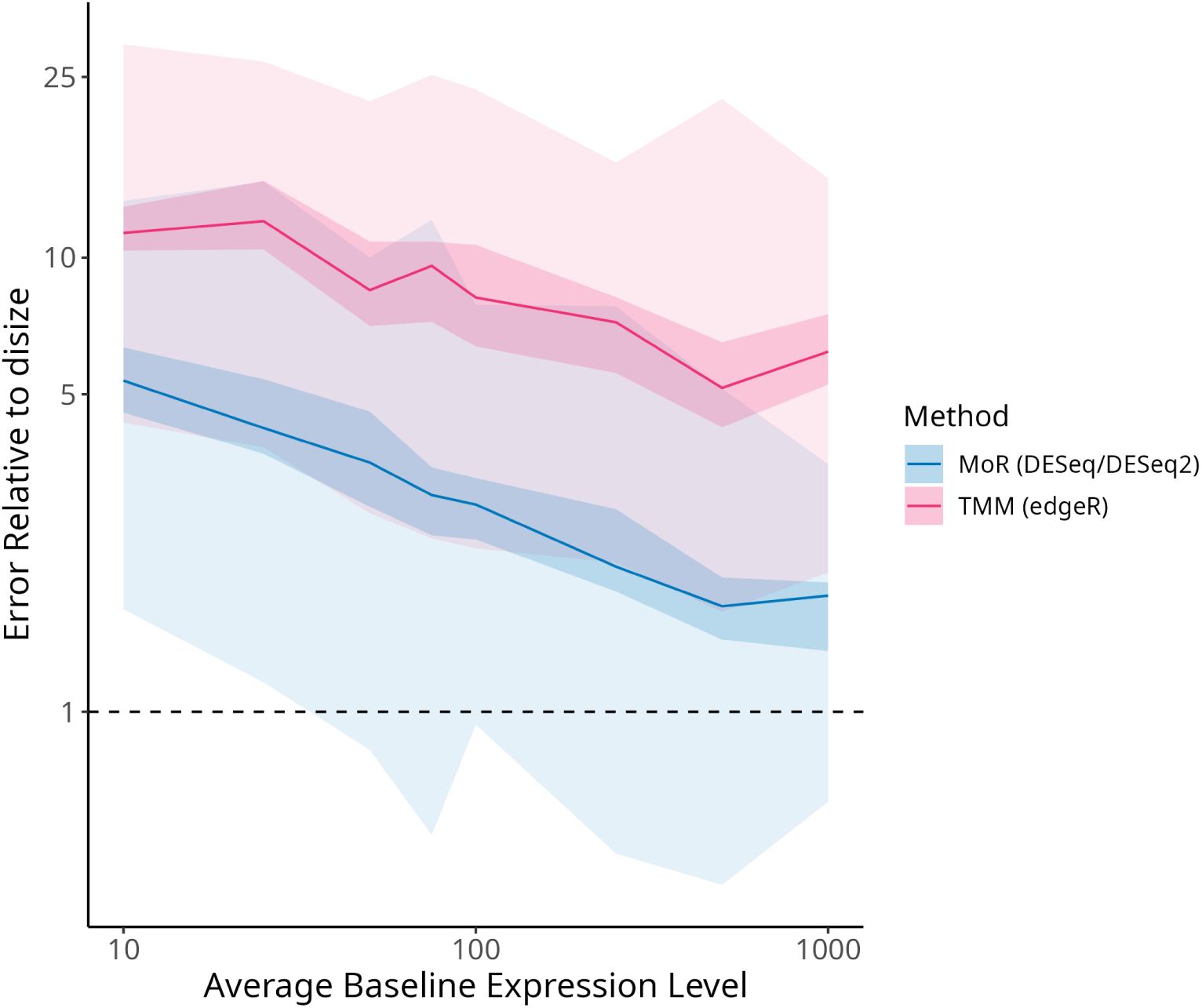
Comparing normalization methods for size factor accuracy across varying expression levels. Fixing *n_g_* = 10000, *p* = 0.2, and *σ* = 2 for the two-condition design, relative error in size factor estimates of MoR and TMM compared to disize versus the average baseline gene expression level. Each line represents the median relative error, while the shaded regions indicate the 5th-95th (lighter) and 40th-60th (darker) percentile ranges. A lower relative error indicates a more accurate size factor estimate.

### Impact On Downstream Analysis

We next assessed the downstream impact of these normalization approaches on differential expression (DE) testing. To emulate a typical analysis workflow, we applied the size factors estimated by each method within the DESeq2 framework. Testing was conducted using the two-condition scenario (10 donors total, 5 replicates per condition) across 100 independent simulation runs (*n*_sims_ = 100). We fixed the true condition effect magnitude at a high variance (*σ* = 2.0) while varying the baseline gene expression levels and testing across highly dense DE environments where 75% to 85% of genes were truly altered by the condition (*p ∈ {*0.15, 0.20, 0.25*}*).

As expected, using the true simulated baseline batch effects yielded the lowest Type I error rates across all baseline expression levels (Fig 2A, S4 Fig), while the Type II error rates remained comparable across all methods (Fig 2B, S5 Fig). The performance of the normalization methods during DE testing directly mirrored their underlying accuracy in size factor estimation, where disize consistently achieved the lowest Type I error compared to MoR and TMM.

**Fig 2.**
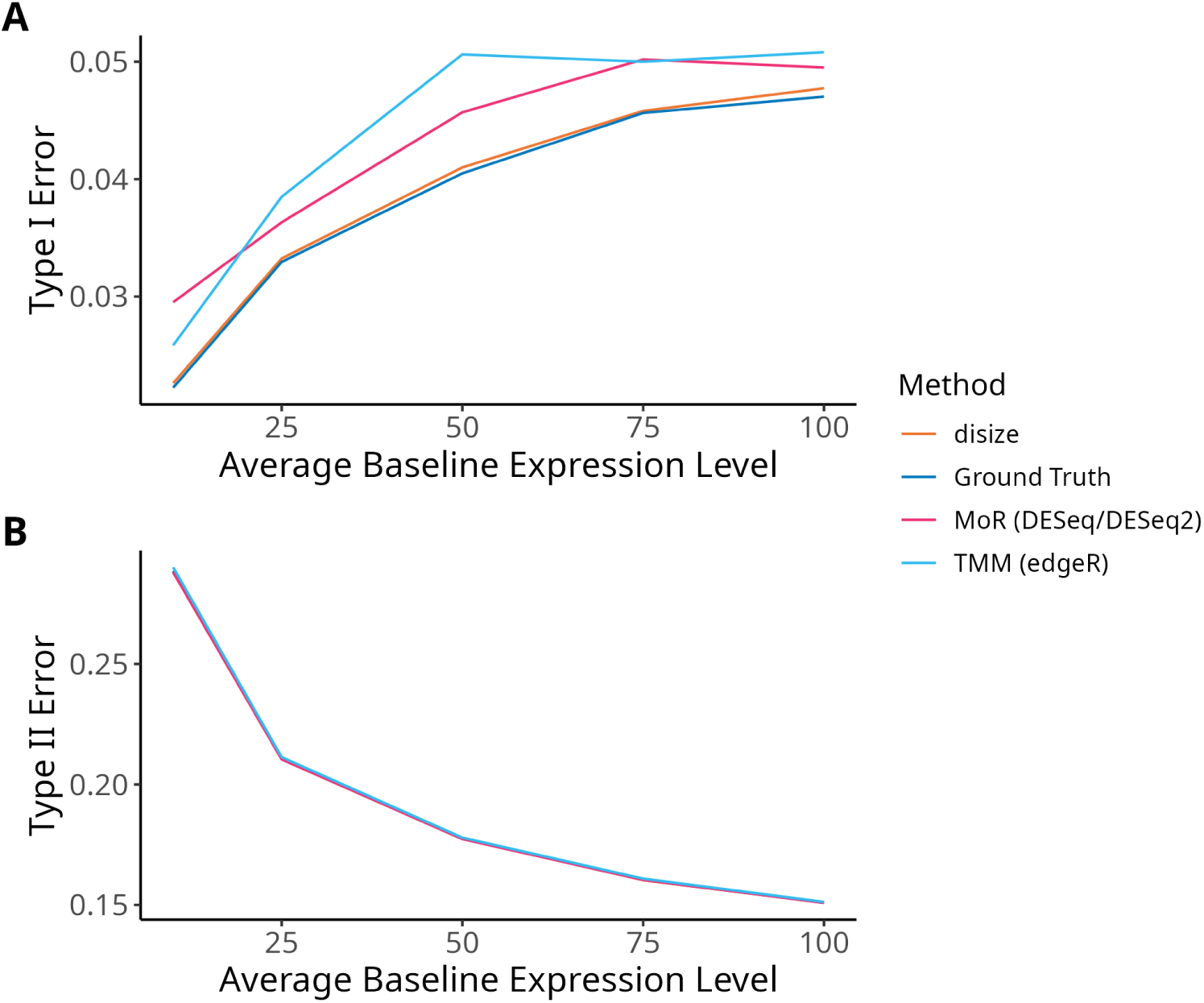
Comparing normalization methods for type I and II error across varying expression levels. Fixing *n_g_* = 10000, *p* = 0.25, and *σ* = 2 under a two-condition design (5 replicates per condition), Type I (A) and Type II (B) error rates of each normalization method (disize, MoR, and TMM) are plotted against the average baseline gene expression level compared to the ground truth. The nominal significance threshold is fixed at *α* = 0.05.

### Validation With Bulk RNA-seq Using SEQC Data

To evaluate disize on bulk transcriptomic data with large-scale expression changes, we used the Sequencing Quality Control (SEQC) consortium benchmark dataset to compare sample A (Universal Human Reference RNA) and sample B (Human Brain Reference RNA) [13]. To replicate typical experiments where spike-ins are absent, we omitted all ERCC features during size factor estimation.

For disize, the experimental design included sample type as a fixed effect, and batch membership was defined as the interaction of sample type, replicate, and sequencing flowcell. We then performed differential expression testing with DESeq2 to evaluate the contrast between samples A and B. Significant differentially expressed genes (DEGs) for each method were identified using a threshold of *p*_adj_ *<* 0.05 and *|*log_2_ fold change*| >* 1. To validate the ability of each method to recover DEGs, we compared these results to independent ground-truth measurements from high-throughput TaqMan quantitative RT-PCR (qRT-PCR) assays, where true DEGs were defined by an absolute expression difference greater than 1. Our benchmark shows that disize is comparable to standard bulk normalization methods, recovering a slightly larger pool of validated DEGs than MoR and TMM (Fig 3).

**Fig 3.**
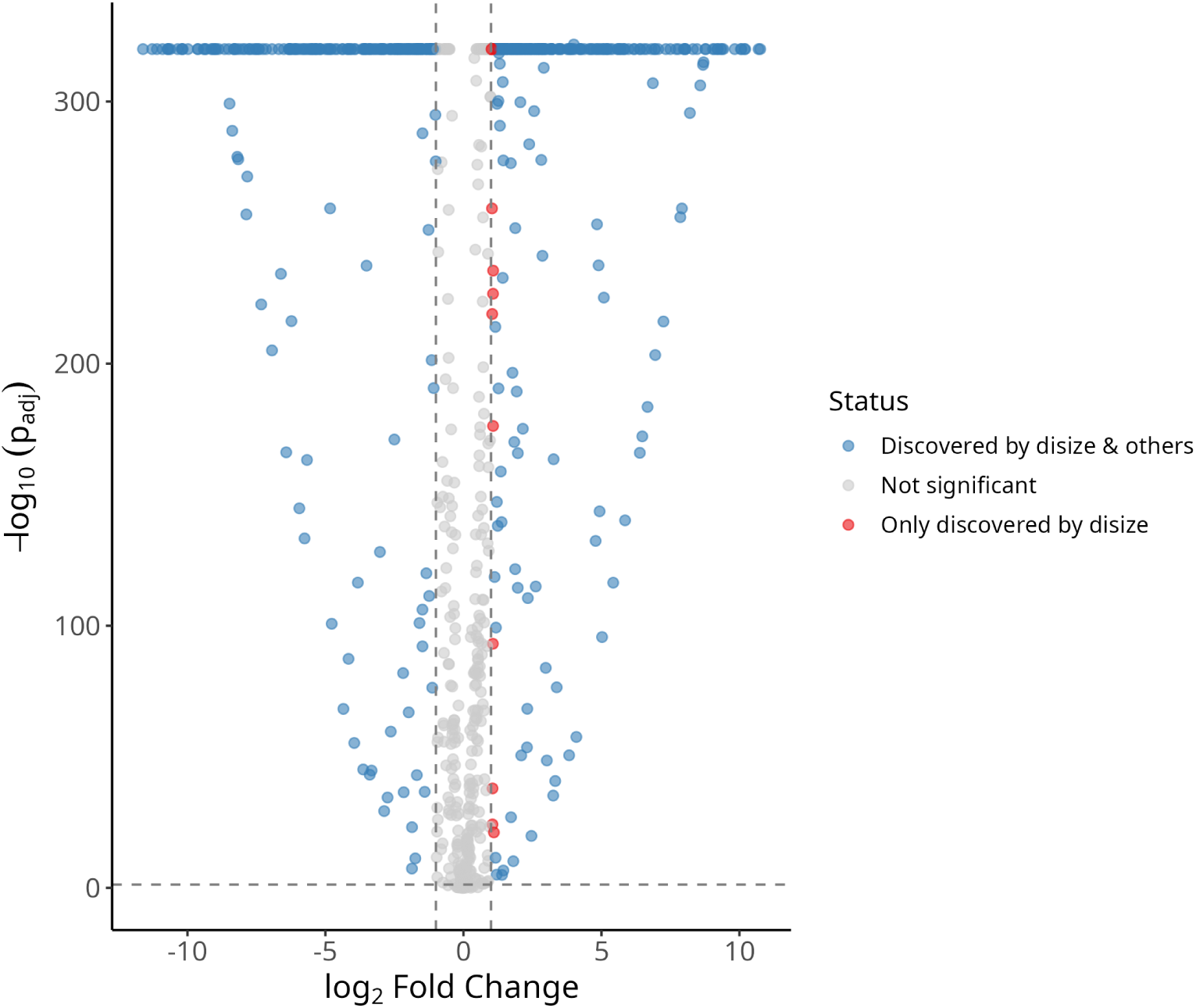
Volcano plot of differential gene expression normalized with disize comparing samples A and B. Points represent individual genes plotted by their log_2_ fold change (B relative to A baseline) and statistical significance (*−* log_10_ *p*_adj_). Genes identified as significant by both disize and alternative normalization methods (MoR/TMM) are shown in blue; genes falling below the significance thresholds (*p*_adj_ *≥* 0.05 or *|*log_2_ fold-change*| ≤* 1.0) are shown in grey; and genes uniquely resolved as significant by disize are shown in red.

### Validation With Pseudo-bulk RNA-seq Data

To evaluate whether the performance advantages of disize persist when confronted with realistic count data, we tested each normalization method on pseudo-bulk data derived from a scRNA-seq dataset. We used a publicly available 10x Genomics dataset consisting of 10,000 Peripheral Blood Mononuclear Cells (PBMCs) [14]. Using the pre-annotated cell-type clusters as distinct biological conditions, we randomly partitioned the cells within each cluster into five batches and performed pseudo-bulking to generate expression profiles for each batch-cluster combination. We next used DESeq2 to perform differential expression analysis on the pseudo-bulk data where no batch effects are present, and then imposed ground-truth size factors on the expression profiles per-batch through binomial thinning. We then benchmarked how well each normalization method recovered the true size factors, the original log_2_ fold-changes, and any genes identified as DE (defined as *p*_adj_ *<* 0.05 and *|*log_2_ fold change*| >* 1).

The pseudo-bulk evaluation closely mirrors the trends observed in our parametric simulations. Both MoR and TMM had higher relative errors in their size factor estimates compared to disize (Fig 4). The error in size factor accuracy directly translated to the accuracy of downstream differential expression testing. When evaluating downstream effects, disize had a lower error in log_2_ fold-change estimates (Fig 5A) as well as a lower Type II error rate (Fig 5B) compared to both MoR and TMM. This confirms that including information from cell-type annotations during normalization allows disize to accurately isolate batch effects in pseudo-bulk data.

**Fig 4.**
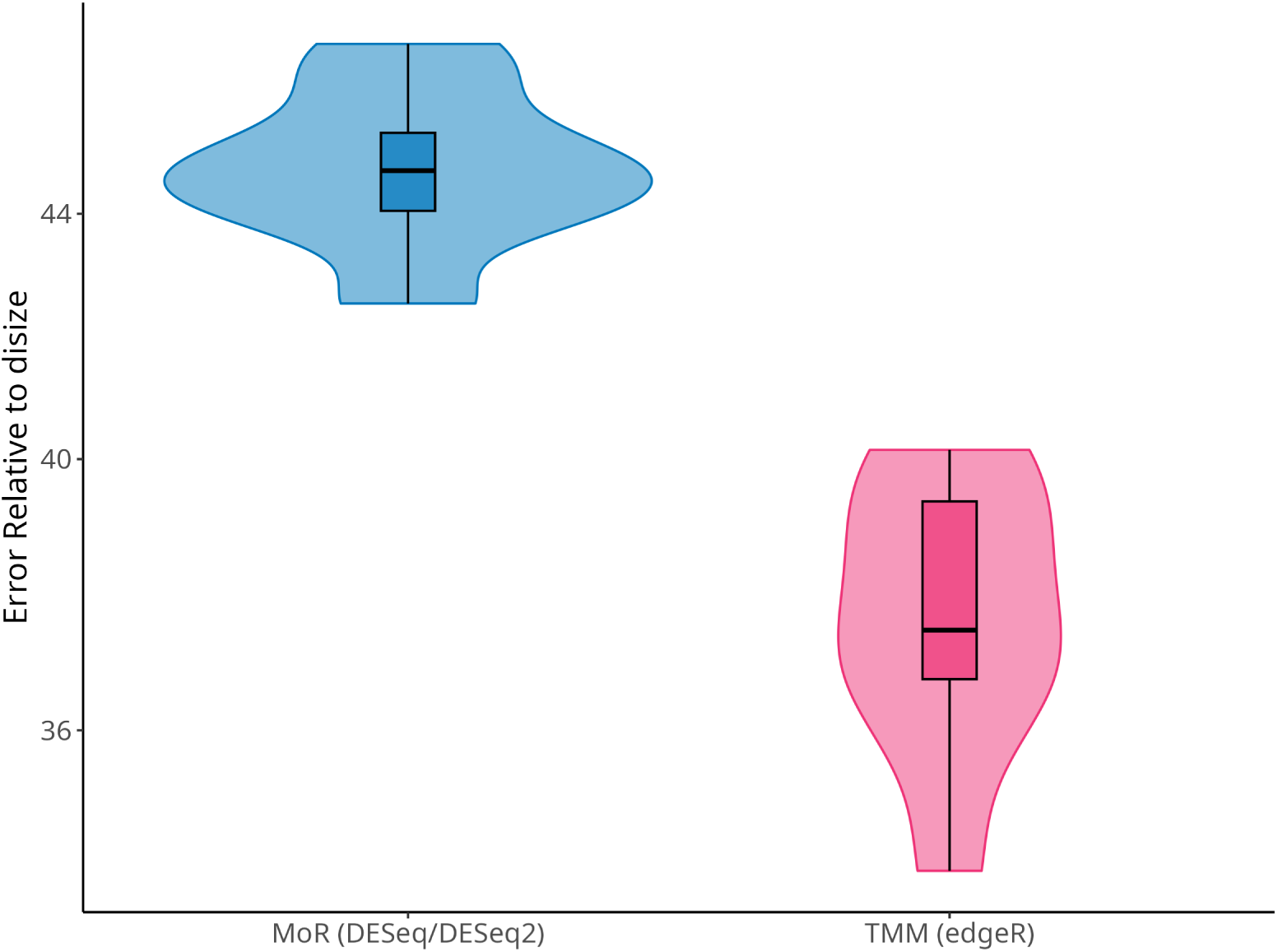
Comparing normalization methods for size factor accuracy on pseudo-bulk data. Benchmarking performance of normalization methods using pseudo-bulk data derived from human PBMC scRNA-seq data. Relative error in size factor estimates of normalization methods compared to disize. A lower relative error indicates a more accurate recovery of the imposed batch effects.

**Fig 5.**
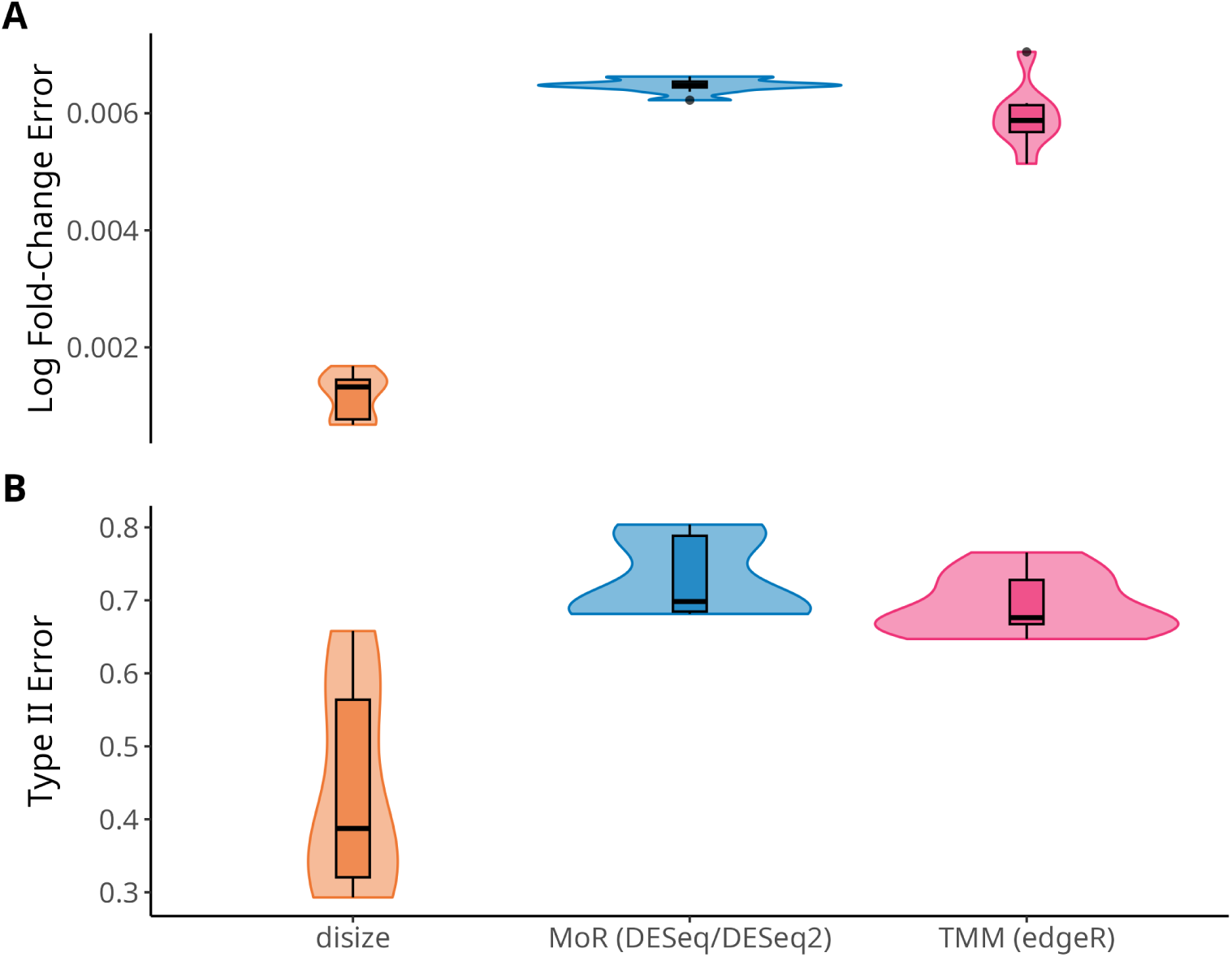
Comparing normalization methods for downstream differential expression analysis on pseudo-bulk data. Benchmarking performance of normalization methods using pseudo-bulk data derived from human PBMC scRNA-seq data. (A) Downstream absolute error in log_2_ fold-change estimates of normalization methods compared to disize in the contrast between clusters 6 and 7. (B) Downstream Type II error of normalization methods compared to disize in the contrast between clusters 6 and 7. A lower error indicates a more accurate recovery of the true differential expression.

### A Case Study of scRNA-seq Data From Interferon-***β*** Stimulated PBMCs

To evaluate its utility on multi-donor single-cell experiments, we applied disize to a scRNA-seq dataset of human PBMCs stimulated with interferon-beta (IFN-*β*) or left untreated as controls [15]. We aggregated the counts into pseudo-bulk profiles stratified by experimental condition, donor identity, and cell-type annotation, yielding a multi-factorial experimental design with a varying number of cells contributing to each pseudo-bulk sample. We then estimated size factors for each sample, where disize was supplied the experimental design with an interaction between the condition and cell-type annotation, and the log-transformed number of cells contributing to each pseudo-bulk sample as offsets.

Using each set of size factors, we conducted cell-type-specific DE testing with DESeq2 by specifying an interaction between the condition and cell-type annotation to compare the IFN-*β* stimulated group against the control baseline within each annotated group. Significant DEGs were identified using a threshold of *p*_adj_ *<* 0.05 and *|*log_2_fold change*| >* 1.

Across the majority of the 13 cell types, disize consistently identified a larger number of significant DEGs compared to MoR and TMM (Fig 6). Notably, this trend was most pronounced in CD14+ monocytes, which yielded the highest absolute number of DEGs discovered by disize (Fig 7). Specifically, disize uniquely resolved the significant downregulation of the oxysterol receptor *GPR183* (EBI2) in CD14+ monocytes. This matches established myeloid feedback mechanisms, where the type I interferon axis and *GPR183* mutually cross-regulate to dictate monocyte activation and trafficking during inflammatory stress [16]. The increased sensitivity relative to other normalization methods indicates that disize is particularly useful for scRNA-seq experiments.

**Fig 6.**
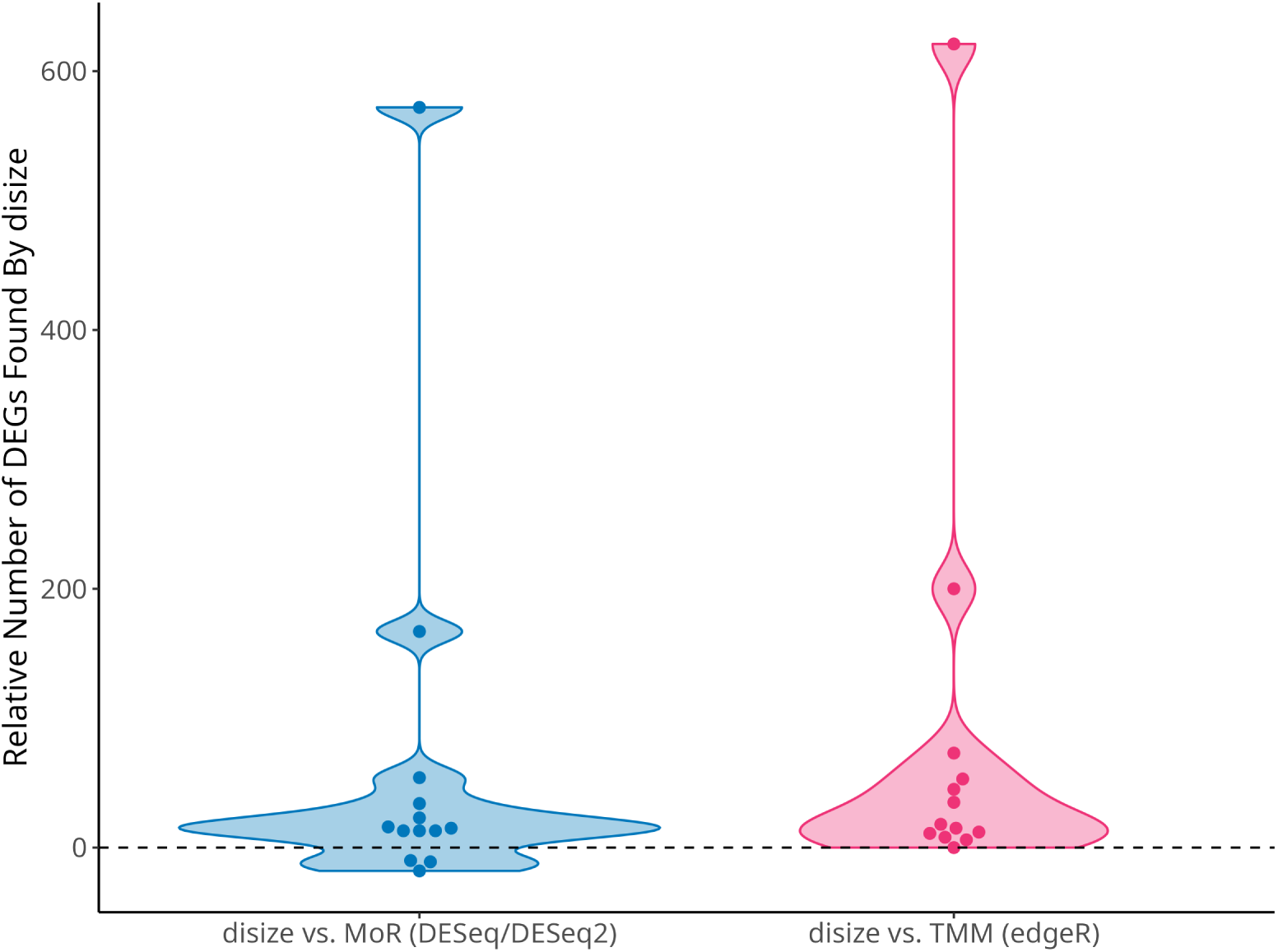
Comparison of identified differentially expressed genes across normalization methods in empirical IFN-*β* stimulated pseudo-bulk data. Relative gain in identified DEGs between conditions (*p*_adj_ *<* 0.05, *|* log_2_ fold-change*| >* 1) for each of the 13 cell types when utilizing disize, calculated as the difference in the number of discovered DEGs compared directly to MoR and TMM. Positive values indicate a higher number of significant genes discovered by disize.

**Fig 7.**
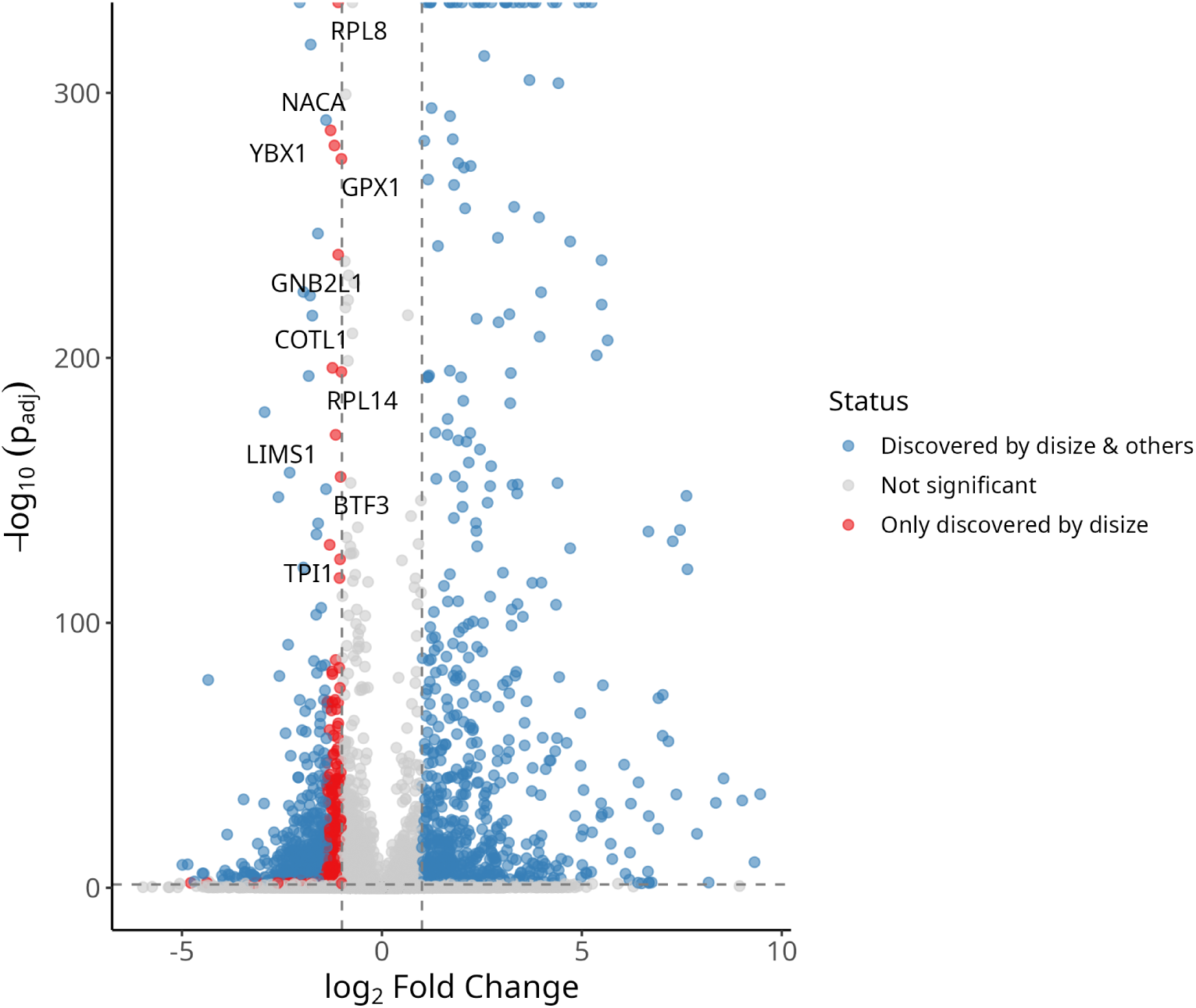
Volcano plot of differential gene expression normalized with disize in CD14+ monocytes. Points represent individual genes plotted by their log_2_ fold change (IFN-*β* stimulated relative to control baseline) and statistical significance (*−* log_10_ *p*_adj_). Genes identified as significant by both disize and alternative normalization methods (MoR/TMM) are shown in blue; genes falling below the significance thresholds (*p*_adj_ *≥* 0.05 or *|*log_2_ fold-change*| ≤* 1.0) are shown in grey; and genes uniquely resolved as significant by disize are shown in red. The top 10 unique disize discoveries by *p*_adj_ are labeled.

## Discussion

While researchers have made significant progress in RNA-seq analysis, much of this work has focused on improving downstream DE testing, with less attention paid to the preceding step of normalization. We address this gap by proposing a novel normalization method that explicitly leverages information from the experimental design to improve the accuracy of size factor estimation.

Our benchmarking demonstrates that disize works well under various scenarios, particularly those with non-trivial biological variation across experimental units, low sparsity (i.e., a high proportion of DE genes), and many genes with low expression. This makes disize particularly well suited for scRNA-seq experiments, where the biological differences between cell types result in little sparsity and pseudo-bulking (aggregating counts from multiple cells into a single “sample”) may not accumulate enough counts for standard methods to work effectively. Indeed, disize greatly outperformed both MoR and TMM in our pseudo-bulk evaluation and recovered more DEGs than MoR and TMM in the case study of IFN-*β* stimulated PBMCs, showing our method successfully leverages information in cell-type annotations to improve size factor estimation.

Although the Type II error rates were comparable across methods in the parametric simulation benchmarks, since disize had the lowest Type I error rate—often below the nominal *α* = 0.05 for low baseline expression—there is potential for increasing the specified significance threshold to produce a lower Type II error rate while retaining a Type I error below the desired *α*. In simpler experiments, however, MoR normalization is roughly as accurate as disize when most genes have sufficient expression (*>* 100 counts) and large systematic biological variation is unlikely.

Previous benchmarks of normalization methods [6, 17] have simulated data by drawing from a multinomial distribution, where the total number of trials represents the library size and a specified vector of probabilities defines the gene expression profile. While this approach is convenient, it fails to account for the overdispersion present in RNA-seq data and lacks a realistic mechanism describing how sample preparation and sequencing imperfectly measure the true number of transcripts resulting in the observed count. We offer an alternative DGP that explicitly models the original magnitude of gene expression and the total effect of technical factors on the observed count distribution.

Since disize jointly estimates the effect of covariates on gene expression and the batch effect, performing inference on the full model may be useful to combine normalization and differential expression quantification into a single step. Currently, we discard the point estimates for the covariate coefficients specified by the experimental design, and do not quantify uncertainty for any parameters. Although the model is complex to fit due to the large number of features in transcriptomic experiments, variational inference methods could approximate the posterior distribution efficiently [18]. This full model would enable differential expression analysis without an explicit normalization step and would quantify uncertainty in size factor estimates.

Although this work has focused on bulk and pseudo-bulk scenarios, since disize allows for an arbitrary batch membership structure, normalization could be done directly on unaggregated scRNA-seq data. This approach leverages the fact that cells belonging to the same sample share a batch effect (avoiding the need for offsets when pseudo-bulking). Consequently, using disize with tools specifically developed for DE testing in single-cell experiments (such as glmGamPoi [19]) a pseudo-bulking step could be avoided altogether. However, an issue with the current implementation is that disize represents the count matrix in a dense format, which increases the memory footprint and may preclude its use with large datasets. Future work could allow a sparse representation, which would significantly reduce the memory requirements.

## Supporting information

**S1 Fig.**
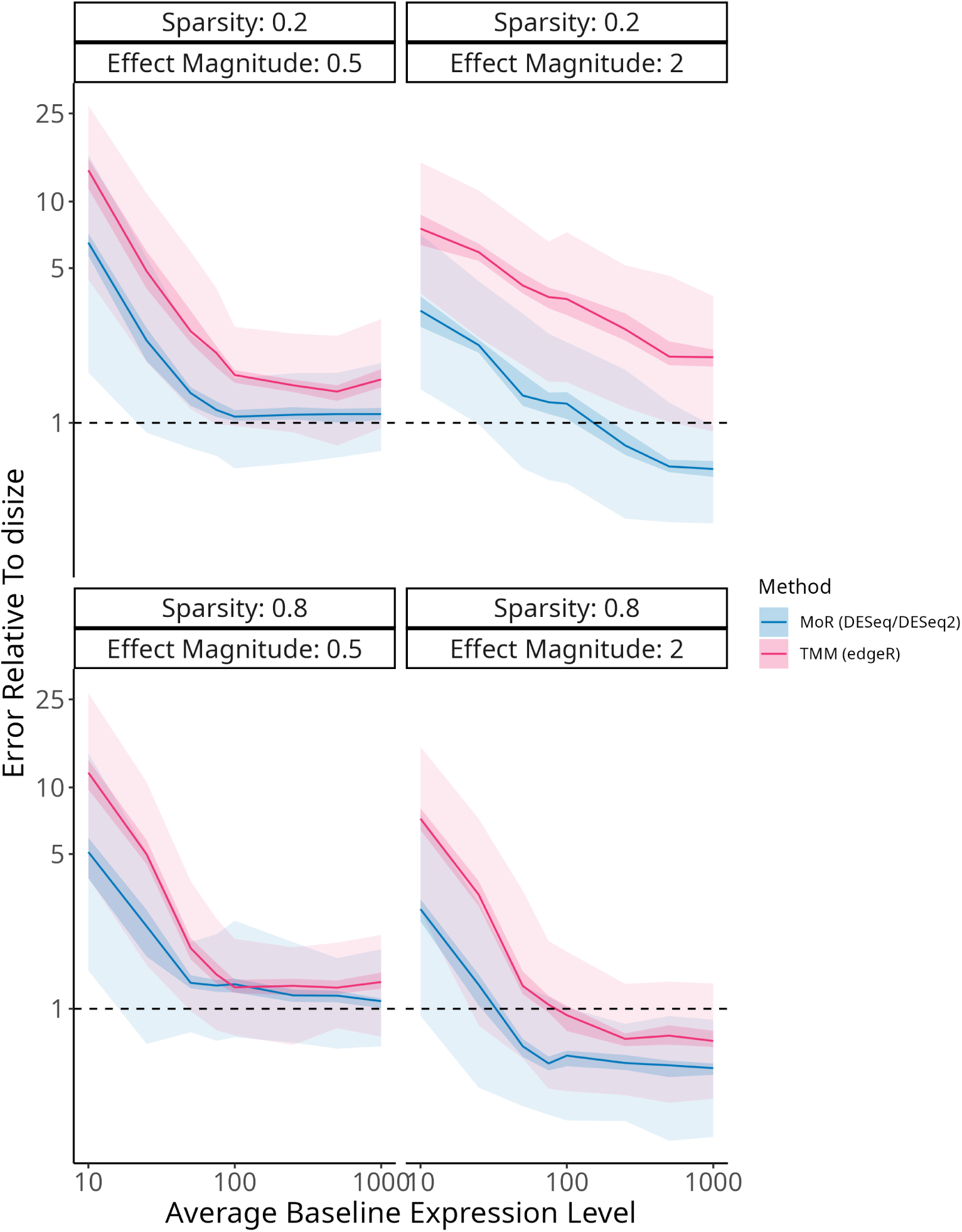
Comparing normalization methods for size factor accuracy in a trivial design. Fixing *n_g_* = 10000 and varying *p* and *σ*, relative size factor error of the other normalization methods (MoR and TMM) compared to disize versus the average baseline gene expression level. Each line represents the median relative error, while the shaded regions indicate the 5th-95th (lighter) and 40th-60th (darker) percentile ranges. A lower relative error indicates a more accurate size factor estimate.

**S2 Fig.**
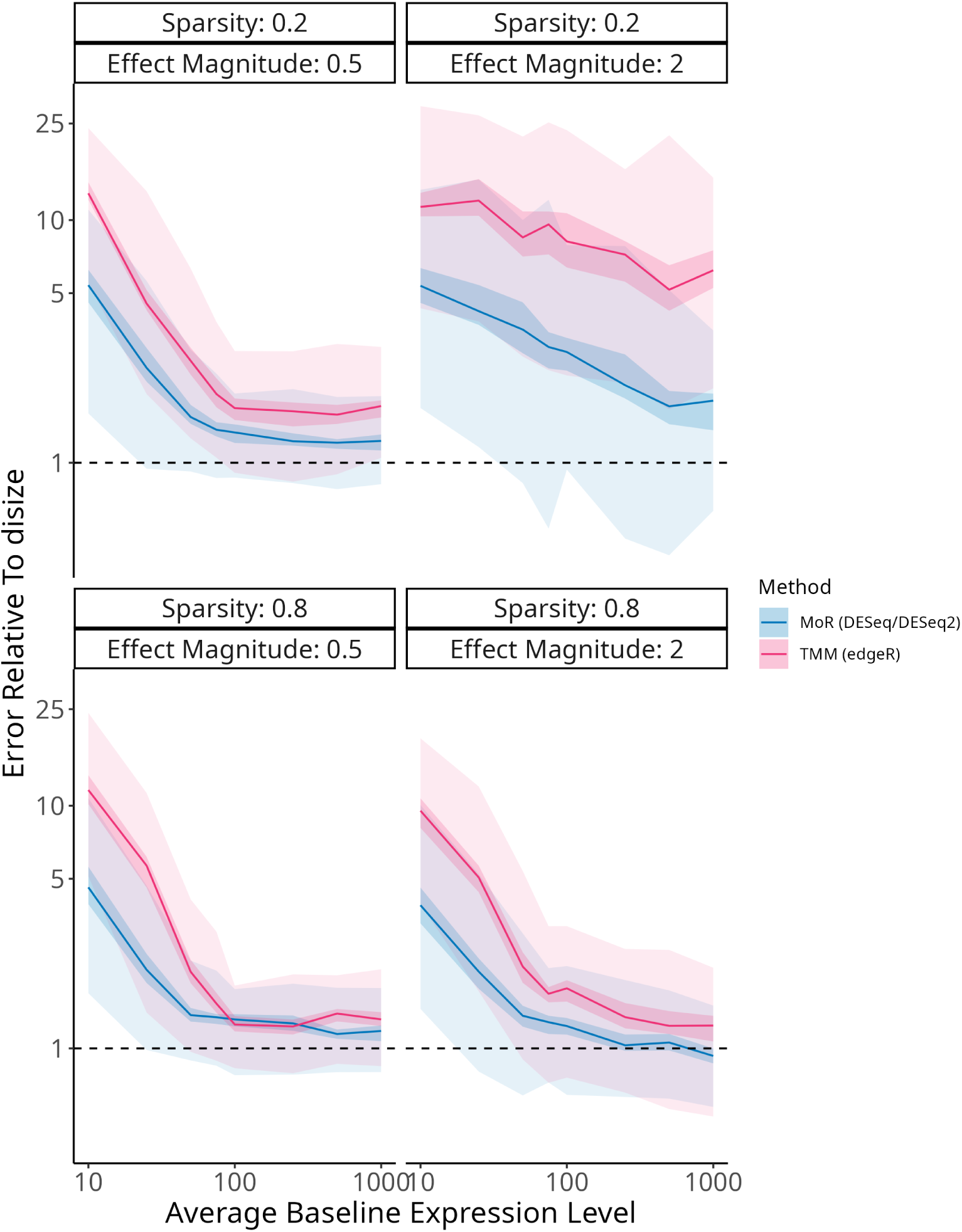
Comparing normalization methods for size factor accuracy in a two-group design. Fixing *n_g_* = 10000 and varying *p* and *σ*, relative size factor error of the other normalization methods (MoR and TMM) compared to disize versus the average baseline gene expression level. Each line represents the median relative error, while the shaded regions indicate the 5th-95th (lighter) and 40th-60th (darker) percentile ranges. A lower relative error indicates a more accurate size factor estimate.

**S3 Fig.**
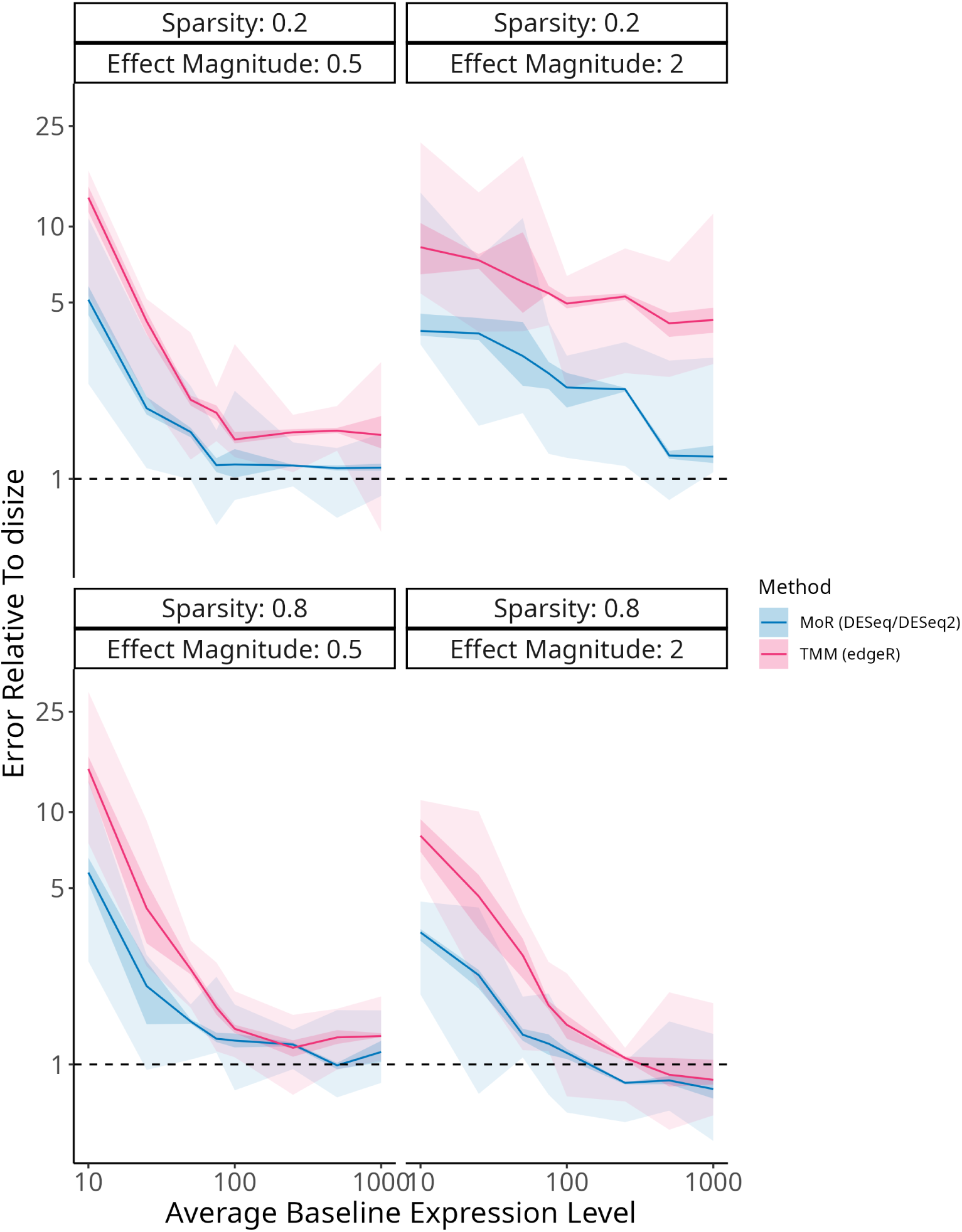
Comparing normalization methods for size factor accuracy in a multifactorial design. Fixing *n_g_* = 10000 and varying *p* and *σ*, relative size factor error of the other normalization methods (MoR and TMM) compared to disize versus the average baseline gene expression level. Each line represents the median relative error, while the shaded regions indicate the 5th-95th (lighter) and 40th-60th (darker) percentile ranges. A lower relative error indicates a more accurate size factor estimate.

**S4 Fig.**
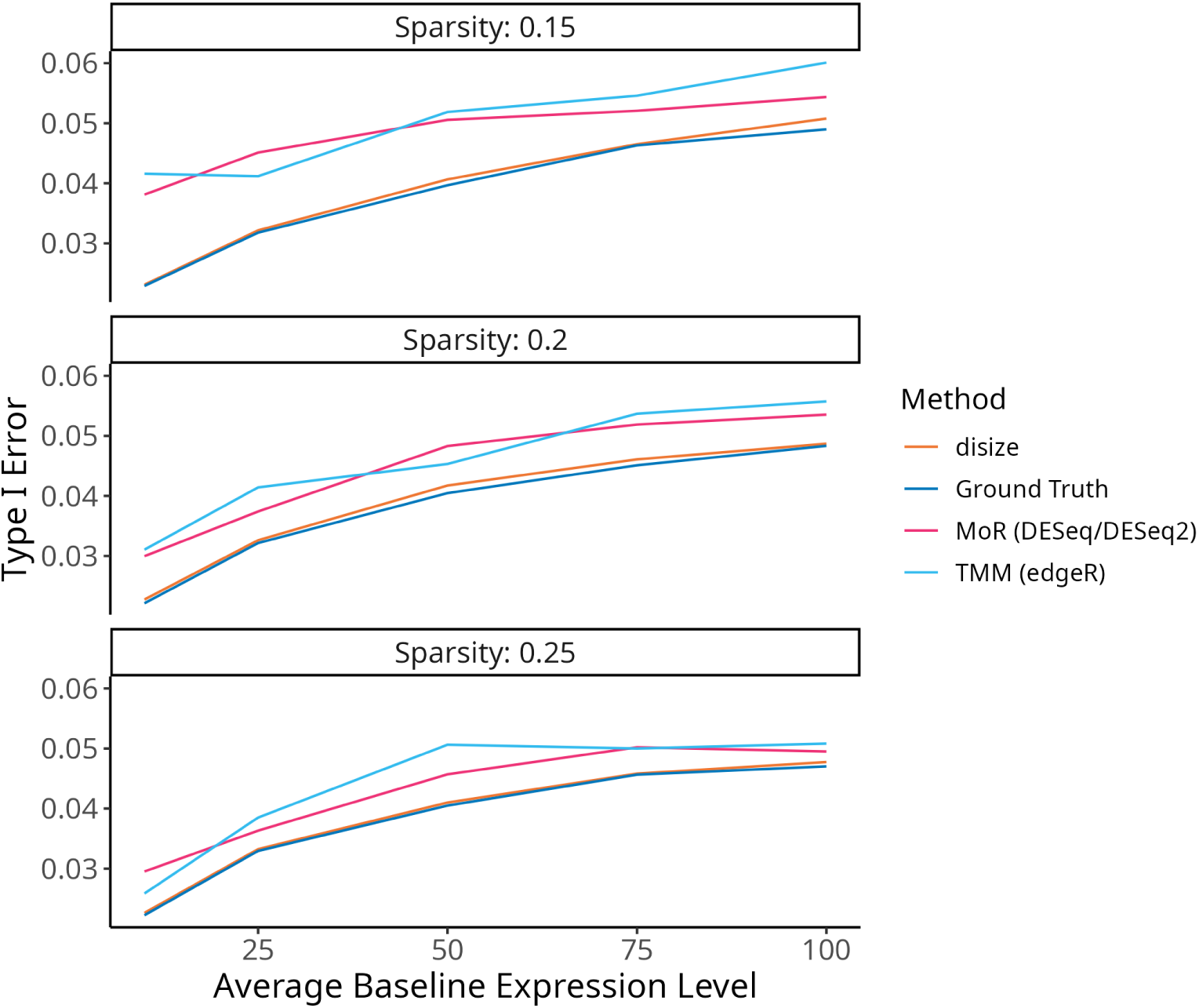
Comparing normalization methods for Type 1 error rates in a two-group design. Fixing *α* = 0.05, *n_g_* = 10000, *σ* = 2, and varying *p*, type 1 error rates of each normalization method (disize, MoR, and TMM) to the ground truth batch effect versus the average baseline gene expression level.

**S5 Fig.**
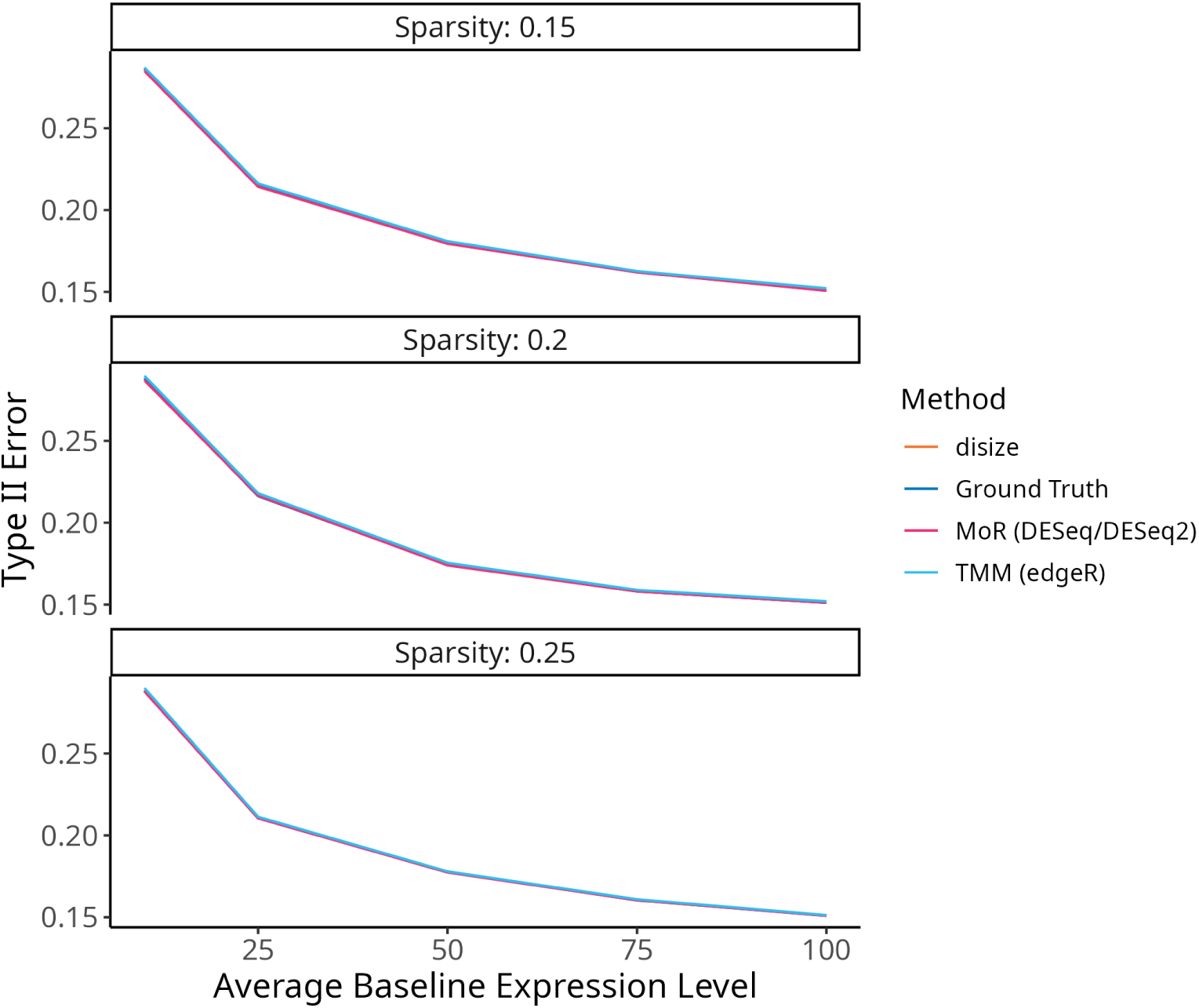
Comparing normalization methods for Type 2 error rates in a two-group design. Fixing *α* = 0.05, *n_g_* = 10000, *σ* = 2, and varying *p*, type 2 error rates of each normalization method (disize, MoR, and TMM) to the ground truth batch effect versus the average baseline gene expression level.

## Notes

### Competing Interest Statement

The authors have declared no competing interest.

